# Variation in Biosynthetic Gene Clusters Among Lifestyles Across Kingdom Fungi

**DOI:** 10.1101/2024.09.14.613087

**Authors:** Brooke M. Allen, Milton Drott, Grant R Nickles, Jason D. Hoeksema

## Abstract

Biosynthetic gene clusters (BGCs) produce secondary metabolites, many of which are involved in trophic interactions and carbon acquisition lifestyles of fungi. As such, BGCs are hypothesized to exhibit evolutionary diversification across those lifestyles. We tested this hypothesis by analyzing predicted BGCs in more than 1000 fungal genomes, and their variation among fungal lineages and across putative fungal lifestyles (trophic modes). Analyses revealed that specific BGC classes, such as PKSI, RiPPs, and terpenes, exhibit significant phylogenetic signals, whereas others, like NRPS and PKS-NRPS Hybrids, do not. BGC abundance was associated with fungal lifestyles: pathotrophs generally had lower BGC abundance, while combined lifestyles such as saprotroph-symbiotrophs had higher BGC abundances. These findings suggest that both phylogenetic history and ecological strategies drive BGC distribution, highlighting their roles in fungal biochemical diversification. Understanding these patterns provides insights into fungal community ecology and the evolutionary processes shaping their biosynthetic potential.

## 1. Introduction

There are an estimated 2-5 million species of fungi on Earth, and since 2010, an average of 1,800 new species have been described each year; however, as of 2017, only around 120,000 species had been formally identified (Blackwell, 2011; Hawksworth and Lücking, 2017). Within this expansive kingdom, fungal species exhibit diverse lifestyles that span across multiple lineages. This complexity challenges the notion that phylogenetic history alone is sufficient for understanding their ecological functions. The study of fungal traits is critical for investigating community assembly and ecosystem function of fungi, as traits associated with resource acquisition, enemy avoidance (such as predation and fungivory), or stress tolerance provide links between the presence and abundance of fungi and their functions and dynamics in communities and ecosystems. Recent reviews highlight measurable and ecologically relevant traits of fungi, including enzymatic activity, mycelial morphology, reproductive structures, dispersion, and habitat colonization (Aguilar-Trigueros et al., 2015; Chaudhary et al., 2022). Symbiotic relationships, in particular, have been noted to influence fungal traits such as spore size and are considered significant drivers in the evolution of fungal characteristics (Aguilar-Trigueros et al., 2023). Understanding these traits is therefore crucial for comprehending fungal biodiversity as well their potential biotechnological applications. However, studying fungal traits presents challenges. Measurements often lack systematic protocols, leading to variability in data collection (Aguilar-Trigueros et al., 2015). Additionally, interpreting traits across different datasets and accounting for high intraspecific variability pose further complexities (Chaudhary et al., 2022). Despite these challenges, advancements in studying fungal traits hold promise for unraveling their roles in ecological processes and applications in various fields. One group of traits found across all fungal lineages and lifestyles, which hold great potential for revealing the ecological roles of fungi, is biosynthetic gene clusters (BGCs) and their encoded chemicals, often referred to as natural products (NPs).

Fungal BGCs and the NPs they produce are increasingly recognized as ecologically relevant traits associated with various functions such as competition, defense mechanisms, and symbiotic relationships (Walsh and Tang, 2017; Keller, 2019; Mudbhari et al., 2023; Zhang et al., 2024). Putative BGCs, which are genetic regions responsible for NP synthesis, are found across diverse taxonomic groups of fungi and can be quantified using various scales including presence/absence, genetic similarity/identity, and broader classifications such as gene cluster families (GCFs) or clans. With the growing availability of fungal genomic data, there is enhanced capability to explore BGC diversity and evolutionary patterns across the fungal kingdom. These chemical profiles are phenotypes shaped by adaptation, linking NP production to the ecology of the fungi that produce them. Investigating how BGCs and associated NP profiles evolve across different fungal lifestyles and lineages can provide valuable insights into fungal community ecology and the ecological functions of NPs (O’Brien and Wright, 2011; Rokas et al., 2018; Naranjo-Ortiz and Gabaldón, 2020). However, although BGCs and NPs are extensively studied in pharmacology for their potential medical applications, there remains a significant gap in understanding their ecological roles. This gap underscores the need for more inclusive research approaches that encompass the full spectrum of fungal biodiversity and lifestyles, thereby advancing our understanding of the ecological roles of BGCs and NPs beyond their pharmacological applications.

Previous studies have indicated that BGCs often exhibit species-specific or narrow taxonomic distributions, with certain fungal lineages or families showing higher productivity of natural products compared to others (Lind et al., 2017; Rokas et al., 2018). Moving beyond taxonomy or evolutionary history, a shift towards functional traits could provide insights into general patterns in fungal ecology (Crowther et al., 2014).

Understanding the ecological roles of fungi involves categorizing them based on trophic modes—such as saprotrophs, pathogens, symbiotrophs, and hybrids—although many species exhibit traits that bridge multiple trophic categories (Zanne et al., 2020). For this study, fungi were broadly categorized into seven lifestyles consisting of saprotroph, pathotroph, symbiotroph and combinations of these three categories. Each lifestyle imposes distinct physiological requirements and selection pressures, which potentially shape the evolution of fungal traits. We aimed to advance understanding of the evolutionary consequences of those selection pressures by investigating how fungal lineage and lifestyle correlate with the abundance and proportion of BGCs across the fungal kingdom. Specifically, we sought to address two main questions: First, whether the proportions and abundances of different BGC classes are associated with specific fungal lifestyles. We hypothesized that variations in resource competition, defense mechanisms, and microbiome interactions among these lifestyles impose natural selection pressures that shape the abundance and proportions of BGC classes.

Second, whether the proportions and abundances of different BGC classes are associated with particular fungal lineages. Here, we hypothesized that closely related fungal lineages will exhibit similar patterns in BGC abundance and proportions, reflecting shared evolutionary histories. By exploring these questions, this study aimed to elucidate whether BGCs can serve as predictive markers of ecological strategies and lineage-specific evolution in fungi, contributing to a deeper understanding of their roles in ecosystem dynamics and evolutionary processes.

## 2. Materials and Methods

We obtained a dataset comprising 1330 fungal genomes along with their annotations (.gff files) from the National Center for Biotechnology Information (NCBI) genome database, selected based on inclusion in the fungal species phylogeny established by Nickles et al. (2023, supplementary table 1). The fungal species tree was generated using 287 gene trees that included single copy orthologs across all species (see details in Nickles et al. (2023)).

To predict putative Biosynthetic Gene Clusters (BGCs), we processed all 1330 fungal genomes using antiSMASH (version 7.1.0) (Medema et al., 2011), with options cb-general, cb-knownclusters, cb-subclusters, pfam2go, and clusterhmmer. Of these, 1281 genomes yielded successful results, with 33 failing to identify BGCs, and 49 facing parsing issues due to gff file formatting inconsistencies (supplementary table 2). The putative BGC classes included in this analysis are: Nonribosomal peptide synthetases (NRPSs), polyketide synthases type I (PKSIs), PKS-NRPS hybrids, ribosomally synthesized and post-translationally modified peptides (RiPPs), terpenes, PKSothers (non-type I PKSs), and others (includes BGCs that do not fit neatly into the previously listed categories).

Trophic mode assignments for the fungal species were determined using FUNGuild via the FUNGuildR package (version 0.2.0.9000) in RStudio (Nguyen et al., 2016). FUNGuild classifies fungal lifestyles into three main categories: pathotrophs, which derive nutrients by harming host cells (including phagotrophs); symbiotrophs, which obtain nutrients through resource exchange with host cells; and saprotrophs, which acquire nutrients by breaking down dead host cells. FUNGuild also includes combinations of these three main categories, reflecting fungi that employ multiple nutritional strategies at some point in their lifecycle. We categorized 1120 of the 1330 genomes into seven generalized FUNGuild trophic modes (symbiotroph, saprotroph-symbiotroph, saprotroph, pathotroph-symbiotroph, pathotroph-saprotroph-symbiotroph, pathotroph-saprotroph, and pathotroph) (supplementary table 3). Of the 1300 genomes, 1009 remained for our analysis after excluding those lacking BGC and/or trophic mode predictions.

For statistical analysis, we employed Bayesian Generalized Linear Mixed Models (GLMM) using the MCMCglmm function from the MCMCglmm package in RStudio (2024.04.1+748 “Chocolate Cosmos”). We treated trophic mode as a fixed effect while genome length and fungal species were included as random effects. Separate models were analyzed for each of the classes of BGCs (PKS, NRPS, terpenes, PKS-NRPS hybrids, RiPPs, Others) and total BGCs, with absolute abundance of the BGC class as the response variable. Phylogenetic and non-phylogenetic versions of models were analyzed; phylogenetic models were fit by including a variance–covariance matrix representing phylogenetic correlations among fungal species, i.e., variance among clades, associated with the random effect for fungal species. Non-phylogenetic models lacked this variance-covariance matrix. The models were run for 10,000 iterations, with a thinning interval of 5 and a burn-in period of 2500. This approach allowed us to investigate the influence of trophic mode on BGC abundance across fungal genomes while controlling for potential confounding factors. Additionally, we compared deviance information criterion (DIC) values between phylogenetic and non-phylogenetic models to provide insights into the presence and significance of phylogenetic signals across different BGC classes. Lower DIC values for phylogenetic models indicated significant phylogenetic signal, indicating meaningful contributions of evolutionary history in explaining variation in genomic abundance of a particular class of BGC.

## 3. Results

Our results suggest that the proportions and abundances of some classes of BGCs are associated with specific fungal lifestyles. For PKSI, pathotrophs exhibited significantly lower abundance compared to pathotroph-saprotroph-symbiotrophs, saprotroph-symbiotrophs, and symbiotrophs, while pathotroph-saprotrophs also showed significantly lower abundance compared to saprotroph-symbiotrophs and symbiotrophs. Additionally, saprotrophs had lower PKSI abundance than saprotroph-symbiotrophs (see Table 1 for statistical details, Figure 1, 2).

**Table 1.**
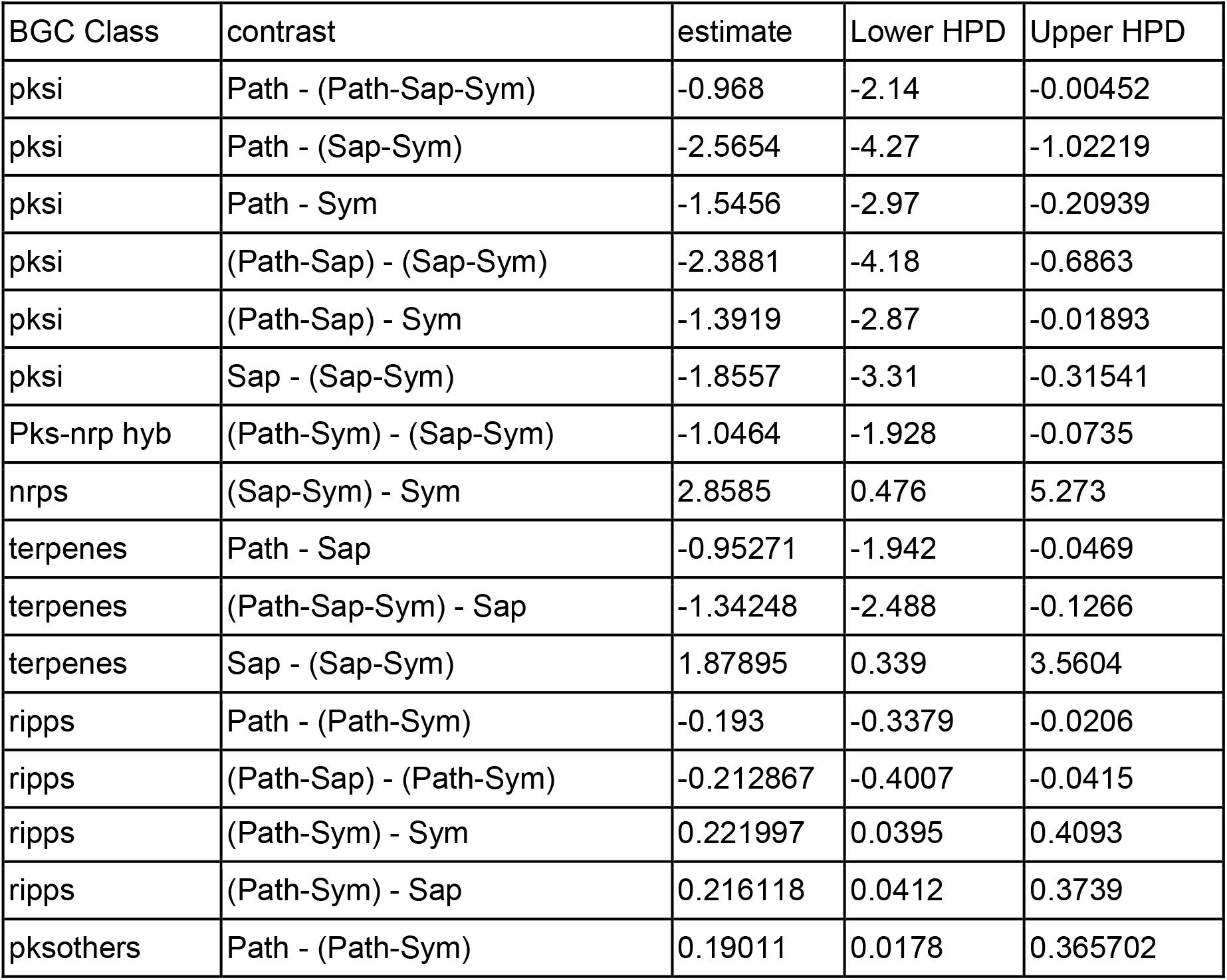
Statistical output for Bayesian Generalized Linear Mixed Models showing significant effects of trophic mode on BGC abundance, accounting for genome length and phylogeny as random effects.

| BGC Class | contrast | estimate | Lower HPD | Upper HPD |
| --- | --- | --- | --- | --- |
| pksi | Path - (Path-Sap-Sym) | -0.968 | -2.14 | -0.00452 |
| pksi | Path - (Sap-Sym) | -2.5654 | -4.27 | -1.02219 |
| pksi | Path - Sym | -1.5456 | -2.97 | -0.20939 |
| pksi | (Path-Sap) - (Sap-Sym) | -2.3881 | -4.18 | -0.6863 |
| pksi | (Path-Sap) - Sym | -1.3919 | -2.87 | -0.01893 |
| pksi | Sap - (Sap-Sym) | -1.8557 | -3.31 | -0.31541 |
| Pks-nrp hyb | (Path-Sym) - (Sap-Sym) | -1.0464 | -1.928 | -0.0735 |
| nrps | (Sap-Sym) - Sym | 2.8585 | 0.476 | 5.273 |
| terpenes | Path - Sap | -0.95271 | -1.942 | -0.0469 |
| terpenes | (Path-Sap-Sym) - Sap | -1.34248 | -2.488 | -0.1266 |
| terpenes | Sap - (Sap-Sym) | 1.87895 | 0.339 | 3.5604 |
| ripps | Path - (Path-Sym) | -0.193 | -0.3379 | -0.0206 |
| ripps | (Path-Sap) - (Path-Sym) | -0.212867 | -0.4007 | -0.0415 |
| ripps | (Path-Sym) - Sym | 0.221997 | 0.0395 | 0.4093 |
| ripps | (Path-Sym) - Sap | 0.216118 | 0.0412 | 0.3739 |
| pksothers | Path - (Path-Sym) | 0.19011 | 0.0178 | 0.365702 |

**Figure 1.**
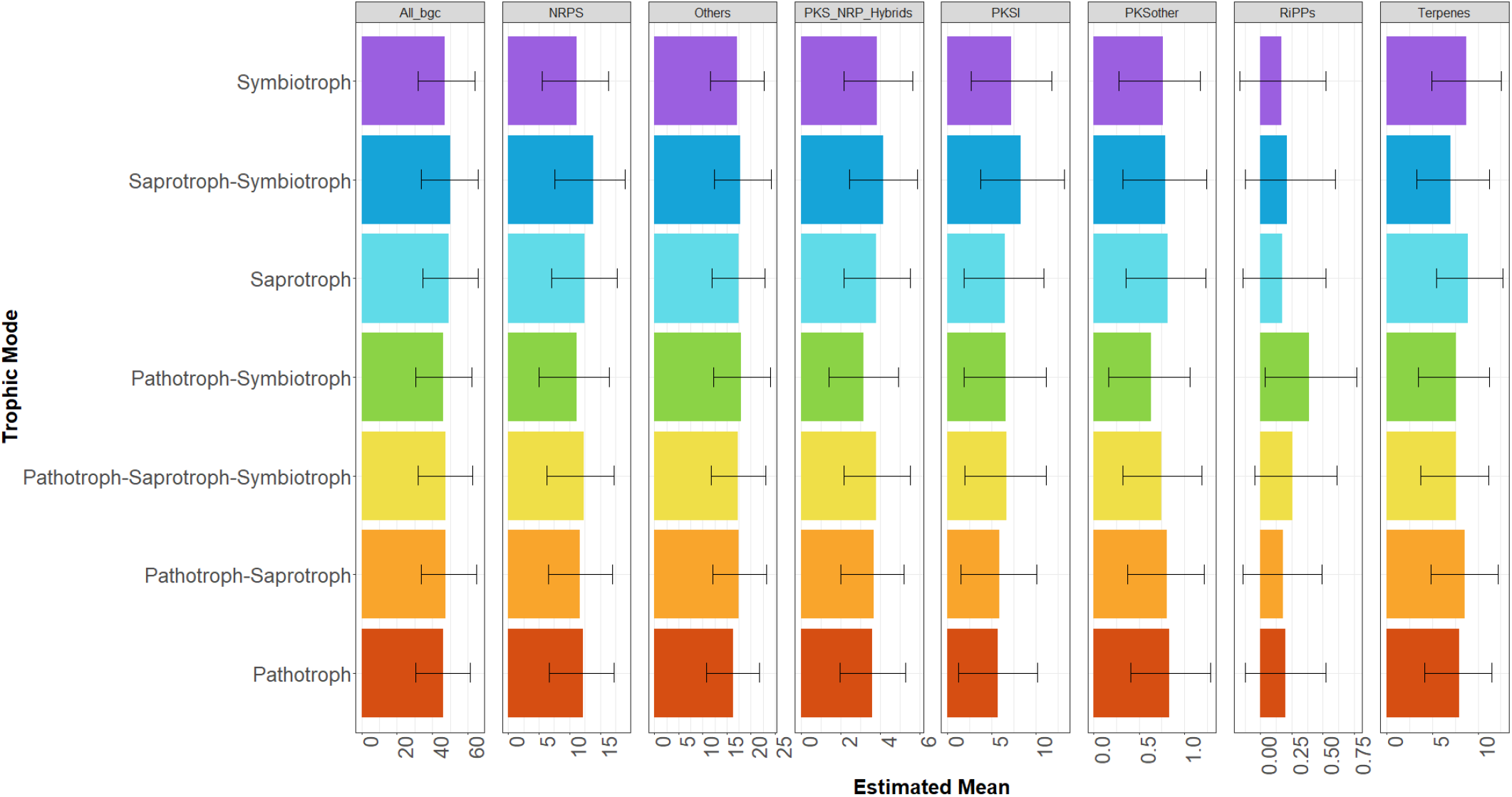
Estimated mean BGC abundance for all BGCs, NRPS, Other, PKS-NRPS hybrids, PKS, PKSothers, RiPPs, and terpenes for each FUNGuild trophic mode.

**Figure 2.**
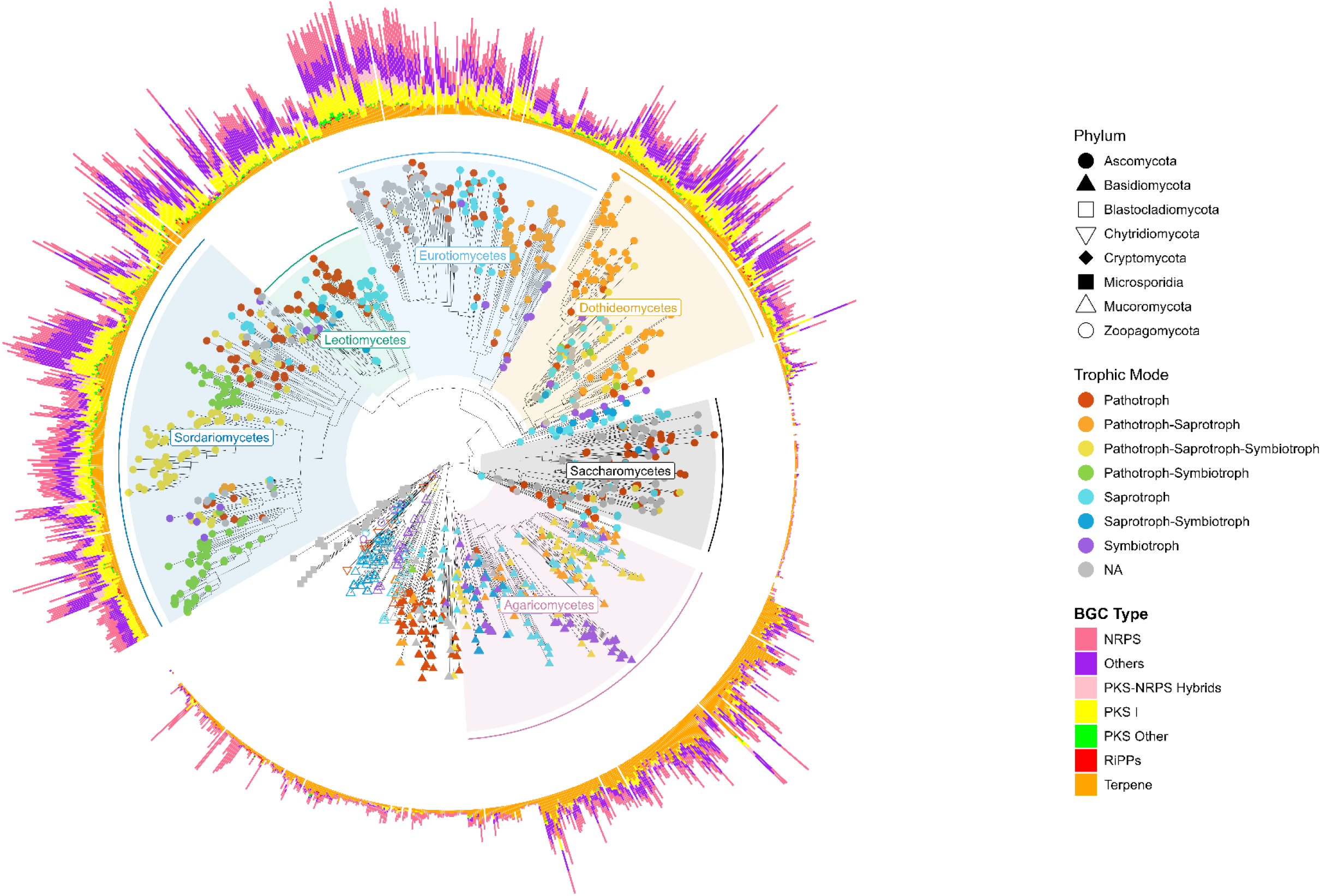
Fungal species tree containing 1330 fungal species. Shapes at the tips represent the phylum of the species while the color of the shapes indicates the FUNGuild trophic mode. The surrounding bar graph shows BGC counts for each of the antiSMASH predicted BGC classes.

For PKS-NRPS hybrids, pathotroph-symbiotrophs had significantly lower abundance than saprotroph-symbiotrophs. For NRPS, saprotroph-symbiotrophs exhibited significantly higher abundance compared to symbiotrophs. For terpenes, pathotrophs had significantly lower abundance than saprotrophs, and pathotroph-saprotroph-symbiotrophs had significantly lower abundance than saprotrophs, whereas saprotrophs had higher abundance compared to saprotroph-symbiotrophs (Table 1, Figure 1, 2).

For RiPPs, pathotrophs had lower abundance compared to pathotroph-symbiotrophs, and pathotroph-saprotrophs showed lower abundance than pathotroph-symbiotrophs. Moreover, pathotroph-symbiotrophs had higher RiPPs abundance compared to both symbiotrophs and saprotrophs. Finally, in the PKSother BGC class, saprotrophs exhibited higher abundance compared to pathotroph-symbiotrophs (Table 1, Figure 1, 2).

Comparison of DIC values for phylogenetic models and non-phylogenetic models revealed that the models for overall BGC abundance, PKS, RiPPs, Terpenes, and Others exhibit a meaningful phylogenetic signal, as indicated by comparatively lower DIC values. This suggests that phylogenetic relationships significantly contribute to explaining the variation in these BGC classes. Conversely, the models for NRPS, PKSother, and PKS-NRPS Hybrids show higher DIC values, indicating that these BGC classes do not exhibit a meaningful phylogenetic signal, and the models without phylogeny fit the data better.

## 4. Discussion

While research on NPs has primarily focused on their pharmaceutical potential, often overlooking their ecological significance, studies have suggested diverse functions for BGCs. These include roles in symbiotic signaling, competition, pathogenicity, and nutrient acquisition among others (Kroken et al., 2003; Lofgren et al., 2024). Our findings support the hypothesis that variations in BGC abundances are associated with specific fungal lifestyles and that closely related fungal lineages exhibit similar patterns in BGC abundance, reflecting shared evolutionary histories. We found that different fungal lifestyles have distinct BGC profiles, and that phylogenetic relationships play a significant role in shaping the distribution and diversity of several BGC classes.

However, some BGC classes, such as NRPS, PKSother, and PKS-NRPS hybrids, did not show meaningful phylogenetic signal, and fungal lifestyles did not exhibit significant differences in Other or overall BGC abundances.

We found that combined trophic modes, such as pathotroph-symbiotroph, often showed higher BGC abundance compared to more strictly defined lifestyles, indicating potential synergistic effects of combining different lifestyles. This trend has been observed previously in specific fungal lineages, such as the family *Xylariaceae*.

Compared to closely related endophytes and saprotrophs in the family *Hypoxylaceae, Xylariaceae* exhibits less distinct ecological modes and increased metabolic diversity (Franco et al., 2022). This versatility in metabolism is thought to result from selection maintaining a diverse array of genes among *Xylariaceae* fungi that blur the lines between saprotrophy and endophytism (Franco et al., 2022). Another known example is *Heterobasidion annosum*, a fungus that is both a forest pathogen and wood decayer, and possesses genes associated with pathogenicity and saprotrophic lifestyles, which are differentially expressed based on the actively engaged lifestyle (Olson et al., 2012). Metabolic diversity is likely beneficial for fungi like *Xylariaceae* and *Heterobasidion annosum* as it enhances their adaptability and survival in various ecological contexts.

Our findings suggest that this trend occurs across the fungal kingdom, indicating that fungi with multiple trophic modes or lifestyles generally have higher BGC abundance for certain types of BGCs.

While BGCs are often associated with competitive or pathogenic interactions (Rokas et al., 2018; Franco et al., 2022), we found that pathotrophs generally had lower abundance across several BGC classes compared to other trophic modes, particularly in PKSI and RiPPs (Figure 1). Natural products produced via BGCs have been identified as virulence factors in numerous fungal pathogens (Howlett, 2006; Macheleidt et al., 2016). For example, the rice blast pathogen *Magnaporthe grisea* utilizes a polyketide, DHN-melanin, as part of its invasion into leaf tissue (Macheleidt et al., 2016). One possibility for the reduced number of PKSI and RiPPs we observed among pathotrophs is that those fungi face trade-offs related to prioritizing the production of specialized virulence factors rather than diversifying their secondary metabolite production. This strategy could enhance their ability to colonize and exploit unique host environments, potentially at the expense of producing a broader range of secondary metabolites.

Previous research has also suggested that pathogen virulence levels evolve to match, but not exceed, the number of resistance genes in the host (Bruns et al., 2014). Given that many fungal pathogens are specialists (Van Der Does and Rep, 2007; Li et al., 2020), this adaptive strategy would allow pathotrophs to optimize their fitness within specific host environments by investing in traits that directly contribute to their pathogenicity and survival. Additionally, having fewer BGCs that are non-essential to fitness might help pathotrophs avoid detection by the host, as there is an ever-present arms race between the host and pathogen (Van Der Does and Rep, 2007). However, the exact mechanisms behind these observations remain unclear, and further research is needed to determine the true causes of this trend.

While the number of BGCs per genomes varies greatly across the fungal kingdom (Robey et al., 2021), we found that more closely related species tend to have similar numbers of BGCs for total BGCs, PKS, RiPPs, Terpenes, and Others. This supports the hypothesis that evolutionary history plays a significant role in shaping BGC diversity and aligns with previous findings that specific BGCs are often narrowly distributed taxonomically (Rokas et al., 2018). However, since we only analyzed total BGC counts within classes and not specific clusters, the conclusions we can draw from this result are limited. Future studies focusing on the analysis of specific BGCs could provide deeper insights into the functional roles and evolutionary patterns of these genomic features across fungal lineages.

Interestingly, we also found a lack of a phylogenetic signal in NRPS, PKSother, and PKS-NRPS Hybrid BGCs, suggesting that closely related species do not have similar abundance for these classes. One potential reason for this lack of phylogenetic signal is horizontal gene transfer (HGT). HGT is a mechanism in fungi and other microbes that facilitates the acquisition of biosynthetic genes from unrelated species, thereby contributing to the non-phylogenetic distribution of these BGCs (Fischbach et al., 2008; Wisecaver and Rokas, 2015; Lind et al., 2017; Rokas et al., 2018). HGT is generally lower in eukaryotic genomes compared to prokaryotic genomes, but there are still many notable examples in fungi (Wisecaver and Rokas, 2015) and it has been speculated that HGT plays a role in the patchy distribution of virulence genes (Van Der Does and Rep, 2007). Another potential contributor to the lack of phylogenetic signal is rapid evolution and turnover of these BGCs, where they are frequently gained, lost, or altered within short evolutionary timeframes, possibly masking any clear phylogenetic signal. This may especially be the case in ‘arms race’ scenarios where fungi rapidly evolve to counteract host defenses or environmental challenges.

This study has several methodological limitations that should be considered. First, genome sampling was limited to a subset of available fungal genomes, which likely do not represent the full diversity of fungal species. Additionally, the FUNGuild tool has limitations in classifying fungi with complex or generalist ecologies and relies on existing knowledge of each species’ ecology for such cases (Nguyen et al., 2016). Finally, the fungal version of AntiSMASH used here (fungiSMASH) was developed primarily with bacteria and Ascomycete non-mycorrhizal fungi (Almeida et al., 2022; Navarro-Munoz and Collemare, 2022), and shows a bias toward known BGC classes with predicted core enzymes, potentially limiting the scope of our results.

Our findings underscore the dual impacts of trophic mode and evolutionary history on BGC abundance in fungal genomes. Moving forward, there is considerable potential to harness these insights through machine learning. Future research directions include developing models that leverage BGC profiles and genomic features to predict the ecological roles of unstudied fungi, identifying potential pathogens, mutualists, or saprotrophs. Integrating phylogenetics with machine learning techniques offers an avenue to predict BGC evolution and distribution across fungal lineages, shedding light on their diversity and ecological functions. Furthermore, applying machine learning to analyze environmental metagenomic data could uncover the ecological roles of fungi within complex microbial communities and their contributions to ecosystem functions.

In conclusion, by taking a trait-based approach we found that both trophic mode and phylogeny significantly influence the abundance of BGCs across the fungal kingdom. Specifically, we observed that pathogens typically exhibit lower levels of PKSI and RiPPs, whereas fungi with combined lifestyles tend to possess higher abundances of PKSI, NRPS, RiPPs, and PKS-NRPS hybrids compared to those with singular lifestyles. Further trait-based approaches focusing on BGCs hold promise for advancing our understanding of fungal ecology and evolution, shedding light on the intricate roles these gene clusters play within ecological settings.

## Supporting information

Supplementary Table 1. List of 1330 NCBI accession numbers

Supplementary Table 2. Antismash BGC predictions

Supplementary Table 3. FUNGuild output

## 6. Acknowledgements

NSF grant #1953299

NSF grant # 1953405

## 7. Supplementary Material

Supplementary Table 1. List of 1330 ncbi accession numbers

Supplementary Table 2. Antismash BGC predictions output

Supplementary Table 3. FUNGuild output

## References

Aguilar-Trigueros, C. A., Hempel, S., Powell, J. R., Anderson, I. C., Antonovics, J., Bergmann, J., et al. (2015). Branching out: Towards a trait-based understanding of fungal ecology. Fungal Biol. Rev. 29, 34–41. doi: 10.1016/j.fbr.2015.03.001

Aguilar-Trigueros, C. A., Krah, F.-S., Cornwell, W. K., Zanne, A. E., Abrego, N., Anderson, I. C., et al. (2023). Symbiotic status alters fungal eco-evolutionary offspring trajectories. Ecol. Lett. n/a. doi: 10.1111/ele.14271

Almeida, H., Tsang, A., and Diallo, A. B. (2022). Improving candidate Biosynthetic Gene Clusters in fungi through reinforcement learning. Bioinformatics 38, 3984–3991. doi: 10.1093/bioinformatics/btac420

Blackwell, M. (2011). The Fungi: 1, 2, 3 … 5.1 million species? Am. J. Bot. 98, 426–438. doi: 10.3732/ajb.1000298

Bruns, E., Carson, M. L., and May, G. (2014). The Jack of All Trades Is Master of None: A Pathogen’s Ability to Infect a Greater Number of Host Genotypes Comes at a Cost of Delayed Reproduction. Evolution 68, 2453–2466. doi: 10.1111/evo.12461

Chaudhary, V. B., Holland, E. P., Charman-Anderson, S., Guzman, A., Bell-Dereske, L., Cheeke, T. E., et al. (2022). What are mycorrhizal traits? Trends Ecol. Evol. 37, 573– 581. doi: 10.1016/j.tree.2022.04.003

Crowther, T. W., Maynard, D. S., Crowther, T. R., Peccia, J., Smith, J. R., and Bradford, M. A. (2014). Untangling the fungal niche: the trait-based approach. Front. Microbiol. 5. doi: 10.3389/fmicb.2014.00579

Fischbach, M. A., Walsh, C. T., and Clardy, J. (2008). The evolution of gene collectives: How natural selection drives chemical innovation. Proc. Natl. Acad. Sci. 105, 4601–4608. doi: 10.1073/pnas.0709132105

Franco, M. E. E., Wisecaver, J. H., Arnold, A. E., Ju, Y.-M., Slot, J. C., Ahrendt, S., et al. (2022). Ecological generalism drives hyperdiversity of secondary metabolite gene clusters in xylarialean endophytes. New Phytol. 233, 1317–1330. doi: 10.1111/nph.17873

Hawksworth, D. L., and Lücking, R. (2017). Fungal Diversity Revisited: 2.2 to 3.8 Million Species. Microbiol. Spectr. 5, 10.1128/microbiolspec.funk-0052–2016. doi: 10.1128/microbiolspec.funk-0052-2016

Howlett, B. J. (2006). Secondary metabolite toxins and nutrition of plant pathogenic fungi. Curr. Opin. Plant Biol. 9, 371–375. doi: 10.1016/j.pbi.2006.05.004

Keller, N. P. (2019). Fungal secondary metabolism: regulation, function and drug discovery. Nat. Rev. Microbiol. 17, 167–180. doi: 10.1038/s41579-018-0121-1

Kroken, S., Glass, N. L., Taylor, J. W., Yoder, O. C., and Turgeon, B. G. (2003). Phylogenomic analysis of type I polyketide synthase genes in pathogenic and saprobic ascomycetes. Proc. Natl. Acad. Sci. 100, 15670–15675. doi: 10.1073/pnas.2532165100

Li, J., Cornelissen, B., and Rep, M. (2020). Host-specificity factors in plant pathogenic fungi. Fungal Genet. Biol. 144, 103447. doi: 10.1016/j.fgb.2020.103447

Lind, A. L., Wisecaver, J. H., Lameiras, C., Wiemann, P., Palmer, J. M., Keller, N. P., et al. (2017). Drivers of genetic diversity in secondary metabolic gene clusters within a fungal species. PLOS Biol. 15, e2003583. doi: 10.1371/journal.pbio.2003583

Lofgren, L., Nguyen, N. H., Kennedy, P. G., Pérez-Pazos, E., Fletcher, J., Liao, H.-L., et al. (2024). Suillus: an emerging model for the study of ectomycorrhizal ecology and evolution. New Phytol. 242, 1448–1475. doi: 10.1111/nph.19700

Macheleidt, J., Mattern, D. J., Fischer, J., Netzker, T., Weber, J., Schroeckh, V., et al. (2016). Regulation and Role of Fungal Secondary Metabolites. Annu. Rev. Genet. 50, 371–392. doi: 10.1146/annurev-genet-120215-035203

Medema, M. H., Blin, K., Cimermancic, P., de Jager, V., Zakrzewski, P., Fischbach, M. A., et al. (2011). antiSMASH: rapid identification, annotation and analysis of secondary metabolite biosynthesis gene clusters in bacterial and fungal genome sequences. Nucleic Acids Res. 39, W339–W346. doi: 10.1093/nar/gkr466

Mudbhari, S., Lofgren, L., Appidi, M. R., Vilgalys, R., Hettich, R. L., and Abraham, P. (2023). Decoding the chemical language of Suillus fungi: genome mining and untargeted metabolomics uncover terpene chemical diversity. bioRxiv, 2023.11.20.567897. doi: 10.1101/2023.11.20.567897

Naranjo-Ortiz, M. A., and Gabaldón, T. (2020). Fungal evolution: cellular, genomic and metabolic complexity. Biol. Rev. 95, 1198–1232. doi: 10.1111/brv.12605

Navarro-Munoz, J., and Collemare, J. (2022). “A bioinformatics workflow for investigating fungal biosynthetic gene clusters,” in Methods in molecular biology (Clifton, N.J.), 1–21. doi: 10.1007/978-1-0716-2273-5_1

Nguyen, N. H., Song, Z., Bates, S. T., Branco, S., Tedersoo, L., Menke, J., et al. (2016). FUNGuild: An open annotation tool for parsing fungal community datasets by ecological guild. Fungal Ecol. 20, 241–248. doi: 10.1016/j.funeco.2015.06.006

Nickles, G. R., Oestereicher, B., Keller, N. P., and Drott, M. T. (2023). Mining for a New Class of Fungal Natural Products: The Evolution, Diversity, and Distribution of Isocyanide Synthase Biosynthetic Gene Clusters. 2023.04.17.537281. doi: 10.1101/2023.04.17.537281

O’Brien, J., and Wright, G. D. (2011). An ecological perspective of microbial secondary metabolism. Curr. Opin. Biotechnol. 22, 552–558. doi: 10.1016/j.copbio.2011.03.010

Olson, Å., Aerts, A., Asiegbu, F., Belbahri, L., Bouzid, O., Broberg, A., et al. (2012). Insight into trade-off between wood decay and parasitism from the genome of a fungal forest pathogen. New Phytol. 194, 1001–1013. doi: 10.1111/j.1469-8137.2012.04128.x

Robey, M. T., Caesar, L. K., Drott, M. T., Keller, N. P., and Kelleher, N. L. (2021). An interpreted atlas of biosynthetic gene clusters from 1,000 fungal genomes. Proc. Natl. Acad. Sci. 118, e2020230118. doi: 10.1073/pnas.2020230118

Rokas, A., Wisecaver, J. H., and Lind, A. L. (2018). The birth, evolution and death of metabolic gene clusters in fungi. Nat. Rev. Microbiol. 16, 731–744. doi: 10.1038/s41579-018-0075-3

Van Der Does, H. C., and Rep, M. (2007). Virulence Genes and the Evolution of Host Specificity in Plant-Pathogenic Fungi. Mol. Plant-Microbe Interactions® 20, 1175–1182. doi: 10.1094/MPMI-20-10-1175

Walsh, C. T., and Tang, Y. (2017). Natural Product Biosynthesis. Royal Society of Chemistry.

Wisecaver, J. H., and Rokas, A. (2015). Fungal metabolic gene clusters—caravans traveling across genomes and environments. Front. Microbiol. 6. doi: 10.3389/fmicb.2015.00161

Zanne, A. E., Abarenkov, K., Afkhami, M. E., Aguilar-Trigueros, C. A., Bates, S., Bhatnagar, J. M., et al. (2020). Fungal functional ecology: bringing a trait-based approach to plant-associated fungi. Biol. Rev. 95, 409–433. doi: 10.1111/brv.12570

Zhang, Z., Zhang, L., Zhang, L., Chu, H., Zhou, J., and Ju, F. (2024). Diversity and distribution of biosynthetic gene clusters in agricultural soil microbiomes. mSystems 9, e01263–23. doi: 10.1128/msystems.01263-23

